# Human nasal and lung tissues infected *ex vivo* with SARS-CoV-2 provide insights into differential tissue-specific and virus-specific innate immune responses in the upper and lower respiratory tract

**DOI:** 10.1101/2021.03.08.434404

**Authors:** Or Alfi, Arkadi Yakirevitch, Ori Wald, Ori Wandel, Uzi Izhar, Esther Oiknine-Djian, Yuval Nevo, Sharona Elgavish, Elad Dagan, Ory Madgar, Gilad Feinmesser, Eli Pikarsky, Michal Bronstein, Olesya Vorontsov, Wayne Jonas, John Ives, Joan Walter, Zichria Zakay-Rones, Menachem Oberbaum, Amos Panet, Dana G. Wolf

## Abstract

The nasal-mucosa constitutes the primary entry site for respiratory viruses including SARS-CoV-2. While the imbalanced innate immune response of end-stage COVID-19 has been extensively studied, the earliest stages of SARS-CoV-2 infection at the mucosal entry site have remained unexplored. Here we employed SARS-CoV-2 and influenza virus infection in native multi-cell-type human nasal turbinate and lung tissues *ex vivo*, coupled with genome-wide transcriptional analysis, to investigate viral susceptibility and early patterns of local-mucosal innate immune response in the authentic milieu of the human respiratory tract. SARS-CoV-2 productively infected the nasal turbinate tissues, predominantly targeting respiratory epithelial cells, with rapid increase in tissue-associated viral sub-genomic mRNA, and secretion of infectious viral progeny. Importantly, SARS-CoV-2 infection triggered robust antiviral and inflammatory innate immune responses in the nasal mucosa. The upregulation of interferon stimulated genes, cytokines and chemokines, related to interferon signaling and immune-cell activation pathways, was broader than that triggered by influenza virus infection. Conversely, lung tissues exhibited a restricted innate immune response to SARS-CoV-2, with a conspicuous lack of type I and III interferon upregulation, contrasting with their vigorous innate immune response to influenza virus. Our findings reveal differential tissue-specific innate immune responses in the upper and lower respiratory tract, that are distinct to SARS-CoV-2. The studies shed light on the role of the nasal-mucosa in active viral transmission and immune defense, implying a window of opportunity for early interventions, whereas the restricted innate immune response in early-SARS-CoV-2-infected lung tissues could underlie the unique uncontrolled late-phase lung damage of advanced COVID-19.

**IMPORTANCE:** In order to reduce the late-phase morbidity and mortality of COVID-19, there is a need to better understand and target the earliest stages of SARS-CoV-2 infection in the human respiratory tract. Here we have studied the initial steps of SARS-CoV-2 infection and the consequent innate immune responses within the natural multicellular complexity of human nasal-mucosal and lung tissues. Comparing the global innate response patterns of nasal and lung tissues, infected in parallel with SARS-CoV-2 and influenza virus, we have revealed distinct virus-host interactions in the upper and lower respiratory tract, which could determine the outcome and unique pathogenesis of SARS-CoV-2 infection. Studies in the nasal-mucosal infection model can be employed to assess the impact of viral evolutionary changes, and evaluate new therapeutic and preventive measures against SARS-CoV-2 and other human respiratory pathogens.

## INTRODUCTION

The ongoing coronavirus disease-2019 (COVID-19) pandemic, caused by severe acute respiratory syndrome coronavirus-2 (SARS-CoV-2), has created an immense global health crisis. While the majority of infections are asymptomatic or cause mild-to-moderate disease, a significant proportion of COVID-19 patients progress over time to display severe pneumonia with acute respiratory distress syndrome, reflecting extensive late-stage viral- and inflammatory-mediated lung injury (1–5). SARS-CoV-2 primarily targets the respiratory tract, utilizing the cellular receptor angiotensin-converting enzyme 2 (ACE-2) and the transmembrane protease serine 2 (TMPRSS2), shown to be expressed in respiratory epithelial cells lining the upper and lower airways (6–13).

The nose is the main port of entry for SARS-CoV-2. The importance of the nasal mucosa as the initial site for SARS-CoV-2 infection is suggested by the observed sequence of clinical manifestations (proceeding from upper-to-lower respiratory involvement), and the higher expression of ACE2 gene in nasal epithelial cells (compared to lower respiratory airway epithelial cells), paralleled by high infectivity of these cells *in vitro* (7, 13).

Frontline protection against respiratory viral infections is mediated by early local-mucosal innate immune responses, exerting antiviral defense via multiple upregulated interferon stimulated genes (ISGs) and cytokines release (14, 15). In the case of SARS-CoV-2, the importance of innate immune defenses in viral control has been highlighted by the finding that inborn defects in innate immunity or the presence of auto-antibodies against interferons are associated with severe COVID-19 (16–18). While the imbalanced innate immune status of end-stage COVID-19, marked by excessive inflammation coupled with impaired interferon production, has been well-characterized (4, 12, 19–21), the earliest innate immune responses to SARS-CoV-2 infection at the nasal mucosal entry site, which could determine the outcome of infection have remained largely unexplored.

Controlled infection studies in animal models, although invaluable for testing vaccines and therapeutics, do not reflect the severe form of the disease in humans (22). Studies of SARS-CoV-2 infection in human airway epithelial cells grown in monolayer cultures and in organoids derived from differentiated lung stem cells have proven instrumental in dissecting the virus biology and cell-type specific interactions (9, 11, 12, 23). However, these models may not recapitulate the tropism of the virus and the host response within authentic multicellular human tissues, that contain a variety of cells of different lineages, as well as extracellular matrix composition - unique to each tissue (24). In this regard, recent work has shown that lung tissue explants can be infected *ex vivo* with SARS-CoV-2, exhibiting impaired interferon (IFN) response with cytokines induction (25, 26).

We have previously reported on the development of *ex vivo* viral infection models in native three-dimensional human target tissues, maintained viable as multi-cell type organ cultures (27–31). We applied these models for the analysis of viral tropism, mode of spread within the tissue, and innate immune effectors of herpes simplex virus type 1, human cytomegalovirus, and Zika virus (32–35). More recently, we have established a novel *ex vivo* model of inferior nasal turbinate tissues – representing the respiratory viral mucosal entry site (36).

In the present study, we employed *ex vivo* SARS-CoV-2 and influenza virus infection in native human nasal turbinate and lung tissues, coupled with genome-wide transcriptional analysis, to investigate viral susceptibility and early patterns of local mucosal defense in the authentic respiratory tract milieu. Our findings provide insights into distinct and virus-specific SARS-CoV-2 -mediated innate immune responses in the upper and lower human respiratory tract.

## RESULTS

### Human nasal turbinate and lung tissues maintained in organ culture are permissive to SARS-CoV-2 infection

The human nasal turbinates are lined by ciliary respiratory epithelium, covering the lamina propria, populated by stromal cells, blood vessels, glands, and immune cells (36). This tissue represents the upper airway entry site for respiratory viruses including SARS-CoV-2. Accordingly, we sought to characterize the susceptibility of the human nasal turbinate tissues, maintained as integral 3D organ cultures, to SARS-CoV-2. We have recently shown that nasal turbinate tissues remain viable and retain their natural histology and functionality, including the continued beating of epithelial cilia, for at least 7 days in culture (36). To further identify tissue-specific and virus-specific aspects of the initial respiratory infection, we have investigated in parallel: 1) SARS-CoV-2 infection in lung tissues, representing the major end-organ site of viral replication and disease [similarly maintained as organ cultures as previously described; (27)], and 2) The susceptibility of the same upper and lower respiratory tract tissues to influenza virus infection.

To evaluate the susceptibility of the nasal turbinate and lung tissues to SARS-CoV-2, we first examined the presence of the SARS-CoV-2 receptor ACE2 and the key protease TMPRSS2, needed for proteolytic cleavage of the viral spike protein. Employing confocal microscopy immunofluorescence analysis of whole-mount tissues, we showed the marked expression of both entry factors in the nasal turbinate tissues, and their colocalization pattern with respiratory epithelial cells lining the mucosa (Figure 1A). In line with latest studies (7, 9, 11, 13), we have also shown the presence of ACE2 and TMPRSS2 proteins in the lung tissues, and their colocalization with epithelial cells lining the alveolar spaces (Figure 1B).

**Figure 1.**
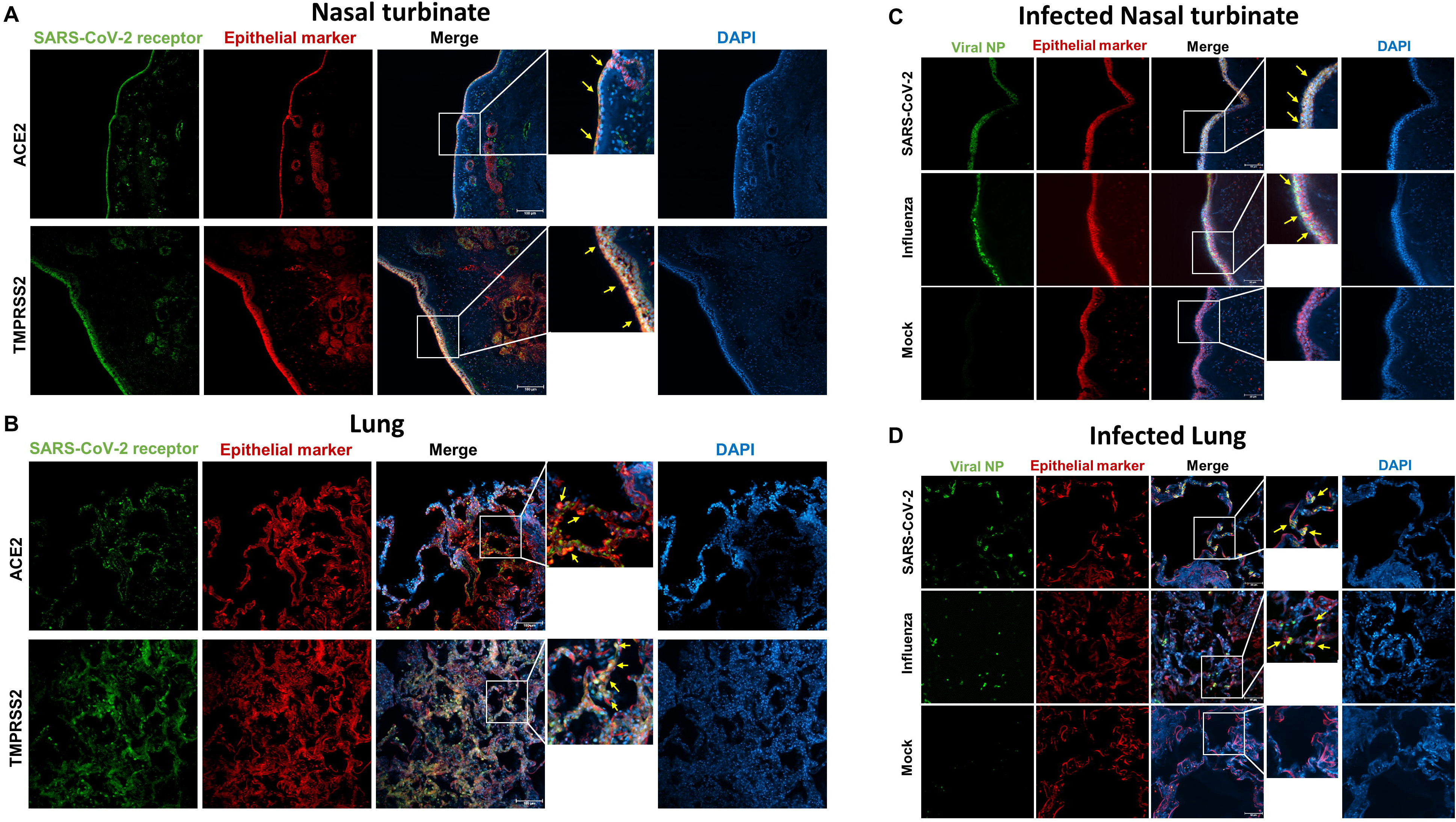
Confocal microscopy analysis of SARS-CoV-2 receptors and cellular tropism in nasal turbinate and lung organ cultures. (**A,B**) Representative confocal micrographs of whole-mount turbinate (**A**) and lung (**B**) cultures, stained for the indicated SARS-CoV-2 receptor and the epithelial cell marker Ep-Cam. Yellow arrows point to cells exhibiting colocalization. DAPI-stained nuclei are shown in blue. Scale bar, 100 μm. (**C,D**) Representative confocal micrographs of whole-mount turbinate (**C**) and lung (**D**) cultures, at 24 hours post infection with SARS-CoV-2 or influenza virus A(H1N1) pdm09 as indicated, showing the respective viral nucleoprotein (NP) colocalization with the epithelial cell marker Ep-Cam. Yellow arrows point to cells exhibiting colocalization. DAPI-stained nuclei are shown in blue. Scale bar, 50 μm.

Next, the turbinate organ cultures were infected with SARS-CoV-2 (isolate USA-WA1/2020). At 2, 24, 48, and 72 hours post infection (hpi), the tissues were washed, and collected along with their respective cleared supernatants. Confocal microcopy analysis of whole-mount infected turbinate tissues at 48 hpi showed the presence of infected cells, primarily localized in the respiratory epithelial cell layer (Figure 1C). No viral immune staining was observed in control mock-infected tissues. We monitored the kinetics of viral infection, measuring the accumulation of the viral sub-genomic (sg) mRNA transcribed in the infected tissues, and the progeny virus genomic RNA released from the infected tissues (measured in the respective cleared supernatants) by quantitative real time (RT)-PCR. As shown in Figure 2A, there was a rapid and significant increase in turbinate-tissue-associated viral sg mRNA (>1- log), and in mature progeny viral RNA released from the infected tissues (∼2.5-log), both reaching peak levels within 24 hpi, followed by a plateau. Consistent with these findings, peak titers (TCID_50_) of newly-formed infectious viral progeny were released from infected tissues at 24 hpi, with declining infectious titers at later times (Figure 2A). Together, these findings revealed productive SARS-CoV-2 replication in the nasal turbinate cultures, with preferential infection of respiratory epithelial cells. Similar viral replication was demonstrated using a low-passage SARS-CoV-2 clinical isolate (SARS-CoV-2 isolate Israel-Jerusalem-854/2020), isolated from a nasopharyngeal swab specimen (data not shown).

**Figure 2.**
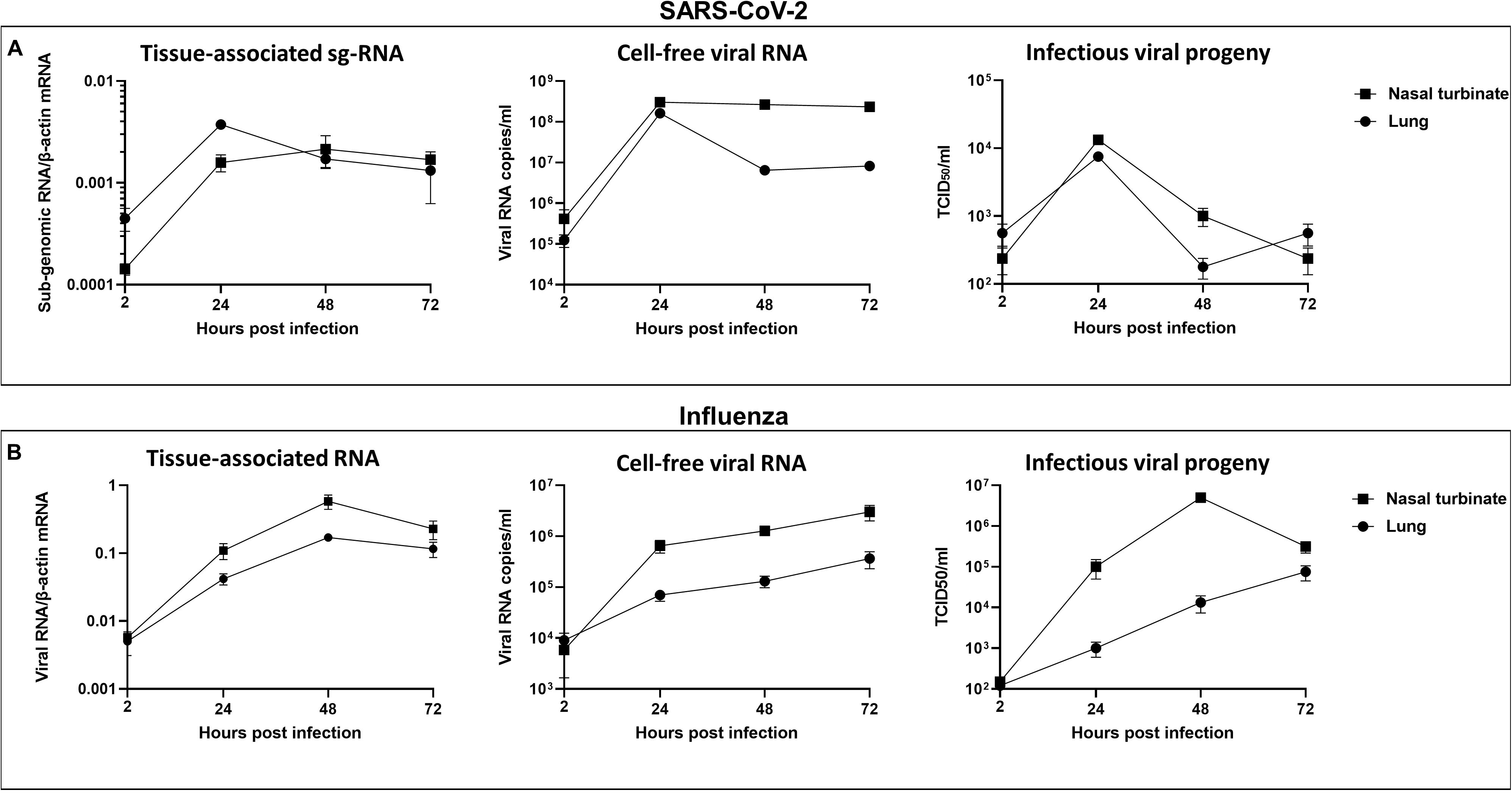
SARS-CoV-2 and Influenza virus infection kinetics in nasal turbinate and lung organ cultures. Nasal turbinate and lung organ cultures were (each) infected in parallel with SARS-CoV-2 (**A**) or influenza A(H1N1) pdm09 virus (**B**) (2×10^5^ TCID_50_/well). (**A**) Levels of tissue-associated SARS-CoV-2 N gene sub-genomic (sg) RNA, determined by qRT-PCR and normalized to β actin (left panel); The copy numbers of extracellular SARS-CoV-2 RNA measured by qRT-PCR in the supernatants of infected tissues (middle panel); Infectious SARS-CoV-2 progeny titers in the supernatants of the same infected tissues, determined by a standard tissue culture infectious dose (TCID)_50_ assay (right panel). (**B**) Levels of tissue-associated Influenza virus RNA, determined by qRT-PCR and normalized to β actin (left panel); The copy numbers of extracellular Influenza virus RNA measured by qRT-PCR in the supernatants of infected tissues (middle panel); Infectious Influenza virus progeny titers in the supernatants of the same infected tissues, determined by a standard TCID_50_ assay (right panel). The data shown are representative of at least 3 independent tissues, and each point represents the mean ± SEM of 5 biological replicates.

In accordance with the presence of viral receptors, we have also demonstrated the active replication of SARS-CoV-2 in *ex vivo* infected lung tissues (Figures 1D, 2A), showing the widespread distribution of the virus in alveolar epithelial cells throughout the tissue. Comparable overall SARS-CoV-2 infection kinetics were observed in the nasal turbinate and in the lung tissues – as measured by viral RNA synthesis and infectious viral progeny levels over time (Figure 2A).

#### Influenza virus efficiently infects the same human nasal turbinate and lung tissues

Influenza virus sialic acid receptors are abundantly expressed in upper and lower respiratory tract epithelial cells (13). To uncover virus-specific patterns of infection and host response, the corresponding tissues from the same donors were infected in parallel with influenza virus A(H1N1) pdm09 (using the same viral inoculum infectious dose). Both the nasal turbinate and the lung tissues were readily infected by influenza virus (Figures 1C,D, 2B). The tissue distribution and respiratory epithelial cell tropism appeared overall comparable to that of SARS-CoV-2, as shown by confocal microscopy analysis (Figure 1C,D). Influenza virus replication and productive infection in the nasal turbinate and lung tissues was demonstrated by the consistent upsurge (∼ 2-log) in tissue-associated influenza RNA and in supernatant cell-free influenza virus genomic RNA levels observed at 24 hpi, with further increase through 48-72 hpi. Unlike the decreasing titers of SARS-CoV-2 noted at 48 and 72 hpi (see above), increasing titers of infectious influenza virus progeny were consistently released from the infected nasal turbinate and lung tissues through these later times post infection (Figure 2B).

### SARS-CoV-2 infection elicits distinct virus-specific and tissue-specific innate immune response patterns in human nasal turbinate and lung tissues

To gain a global insight into the earliest tissue responses to SARS-CoV-2, we employed unbiased genome-wide transcriptome analysis of infected versus mock-infected tissues at 24 hpi. This time point was chosen based on our demonstration that SARS-CoV-2 replication in the tissues already reaches its peak at 24 hpi (see Figure 2), and our preliminary RT-PCR analysis of individual cytokines kinetic in SARS-CoV-2- and influenza virus -infected nasal turbinate tissues (data not shown), indicating that the transcriptional response to SARS-CoV-2 and influenza virus in infected tissues is well induced by 24 hpi. We sought to define common versus virus-specific and tissue-specific innate immune response patterns, by comparing the transcriptional response of the nasal turbinate and lung tissues to SARS-CoV-2 and influenza A(H1N1) pdm09 (upon parallel infection of the same tissues, as described above).

Transcriptome analysis was carried out independently on 3 nasal tissues and 5 lung tissues (all obtained from different donors) to gain statistical significance. We detected viral gene transcripts representing coverage over the entire viral genome in all infected tissues (data not shown). The percent of viral transcripts (of all the sequence reads in the transcriptome data) was overall comparable upon infection by SARS-CoV-2 and influenza virus in both tissues, except for a significantly higher percent of SARS-CoV-2 transcripts reads in the lung tissues compared to the percent of SARS-CoV-2 reads in the nasal turbinate tissues (Figure S1).

#### Innate responses of the infected nasal turbinate tissues

To start evaluating the global host response pertaining to each of the conditions, the samples were grouped in principal-component analysis (PCA). As shown, SARS-CoV-2 and influenza exerted distinct global signatures in the nasal turbinate tissues, reflected by their disparate coordinates (Figure 3A); SARS-CoV-2 and influenza transcriptional signatures differentially distributed from mock along PC2, with more progressive transcriptional response observed along this axis in SARS-CoV-2 infected tissues. We found that SARS-CoV-2 infection substantially affected the global gene expression profile in the nasal turbinate tissues, leading to differential expression of 371 genes (309 upregulated and 62 downregulated following infection; Figure 3B). The most profoundly upregulated genes included antiviral ISGs, cytokines and chemokines (Figures 3C, S2A). Of note, in addition to familiar ISGs, one of the most upregulated genes in SARS-CoV-2 infected nasal turbinate tissues was the long non-protein coding RNA LINC00487 (Figure 3C), recently identified as a novel ISG (37). Employing qRT-PCR of RNA purified from independent infected and mock-control nasal turbinate tissues, we validated the viral-induced upregulation of selected innate immunity genes following infection (Figure S3). In accordance, and further defining the affected biological pathways and predicted functions, Ingenuity Pathway Analysis (IPA) showed that SARS-CoV-2 infection in the nasal mucosal tissues primarily induced antiviral and proinflammatory pathways related to interferon signaling, innate immunity, and immune cell activation (Figure 3C,D, see also Figure 5A). Despite similar levels of infection, we observed some transcriptional response variability between the three independent nasal turbinate tissues (mainly with respect to the extent, but not the direction: up-versus downregulation, of differential gene response; Figure 3D). As we have shown before, this tissue-to-tissues variability, reflecting the natural diversity between individuals, is expected in studies involving human tissues (36).

**Figure 3.**
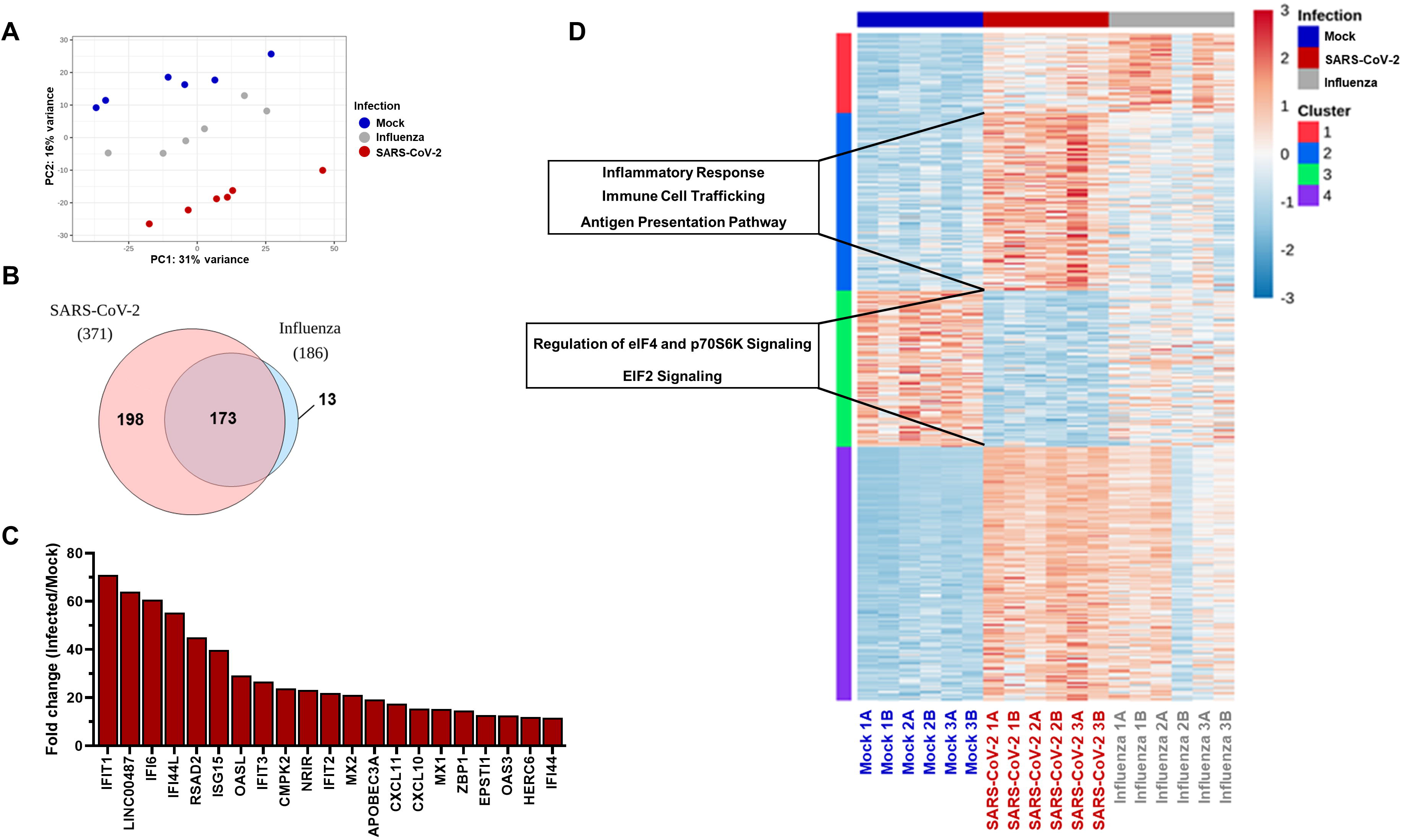
Nasal turbinate tissue transcriptional response to SARS-CoV-2 and Influenza virus infection. Nasal turbinate tissues were mock-infected or infected in parallel with SARS-CoV-2 or influenza A(H1N1) pdm09 (2×10^5^ TCID_50_/well). At 24 hours post infection, RNA was extracted and subjected to transcriptome analysis. Three independent donors’ tissues (with two biological replicates for each experimental condition) were included in the analysis. (**A**) Principal-component analysis (PCA) of the global transcriptional response of the nasal turbinate tissues to SARS-CoV-2 or Influenza virus infection. The first two PCs are shown. (**B**) Venn diagram illustration of the number of unique and overlapping differentially expressed (DE) genes in SARS-CoV-2 and Influenza virus infected nasal turbinate tissues. (**C**) The twenty most-profoundly upregulated genes in SARS-CoV-2-infected (versus mock-infected) nasal turbinate tissues. (**D**) Clustered heatmap representation of all genes with a significant contribution of infection to their expression in nasal turbinate tissues. Normalized expression values were scaled at gene level (scale is shown at top-right), then clustered by kmeans (with k=4), as indicated. Representative pathways and molecular functions distinctively enriched in SARS-CoV-2 infected tissues (versus influenza infected tissues), as reflected by the related upregulated genes in cluster 2 and downregulated genes in cluster 3, are indicated at the left.

**Figure 4.**
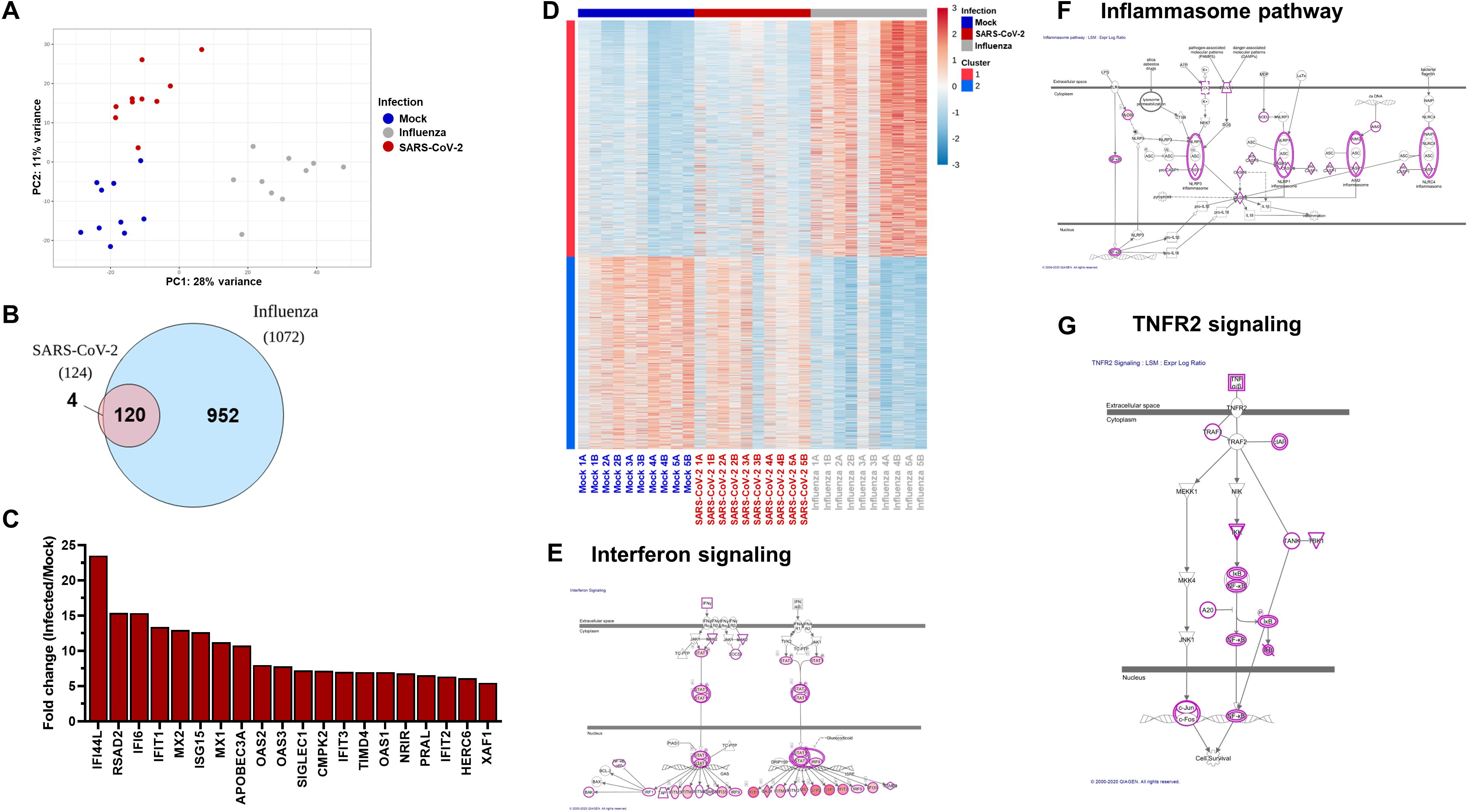
Lung tissue transcriptional response to SARS-CoV-2 and Influenza virus infection. Lung tissues were mock-infected or infected in parallel with SARS-CoV-2 or influenza A(H1N1) pdm09 (2×10^5^ TCID_50_/well). At 24 hours post infection, RNA was extracted and subjected to transcriptome analysis. Five independent donors’ tissues (with two biological replicates for each experimental condition) were included in the analysis. (**A**) Principal-component analysis (PCA) of the global transcriptional response of the lung tissues to SARS-CoV-2 or Influenza virus infection. The first two PCs are shown. (**B**) Venn diagram illustration of the number of unique and overlapping differentially expressed (DE) genes in SARS-CoV-2 and Influenza virus infected lung tissues. (**C**) The twenty most-profoundly upregulated genes in SARS-CoV-2-infected (versus mock-infected) lung tissues. (**D**) Clustered heatmap representation of all genes with a significant contribution of infection to their expression in lung tissues. Normalized expression values were scaled at gene level (scale is shown at top-right), then clustered by kmeans (with k=2), as indicated. (**E-G**) IPA® overlapping schemes of Interferon Signalling (**E**), Inflammasome (**F**) and TNFR2 Signalling (**G**) canonical pathways. Significantly differentially expressed genes between SARS-CoV-2 infected and mock-infected tissues are overlaid with those that were identified when comparing Influenza infected to mock-infected tissues (encircled by a pink line). Upregulated genes are coloured in shades of red from white (not significantly changed), to dark red (highly upregulated). Empty pink circles stand for differentially expressed genes that were found in lung tissues infected with Influenza virus but not with SARS-CoV-2.

**Figure 5.**
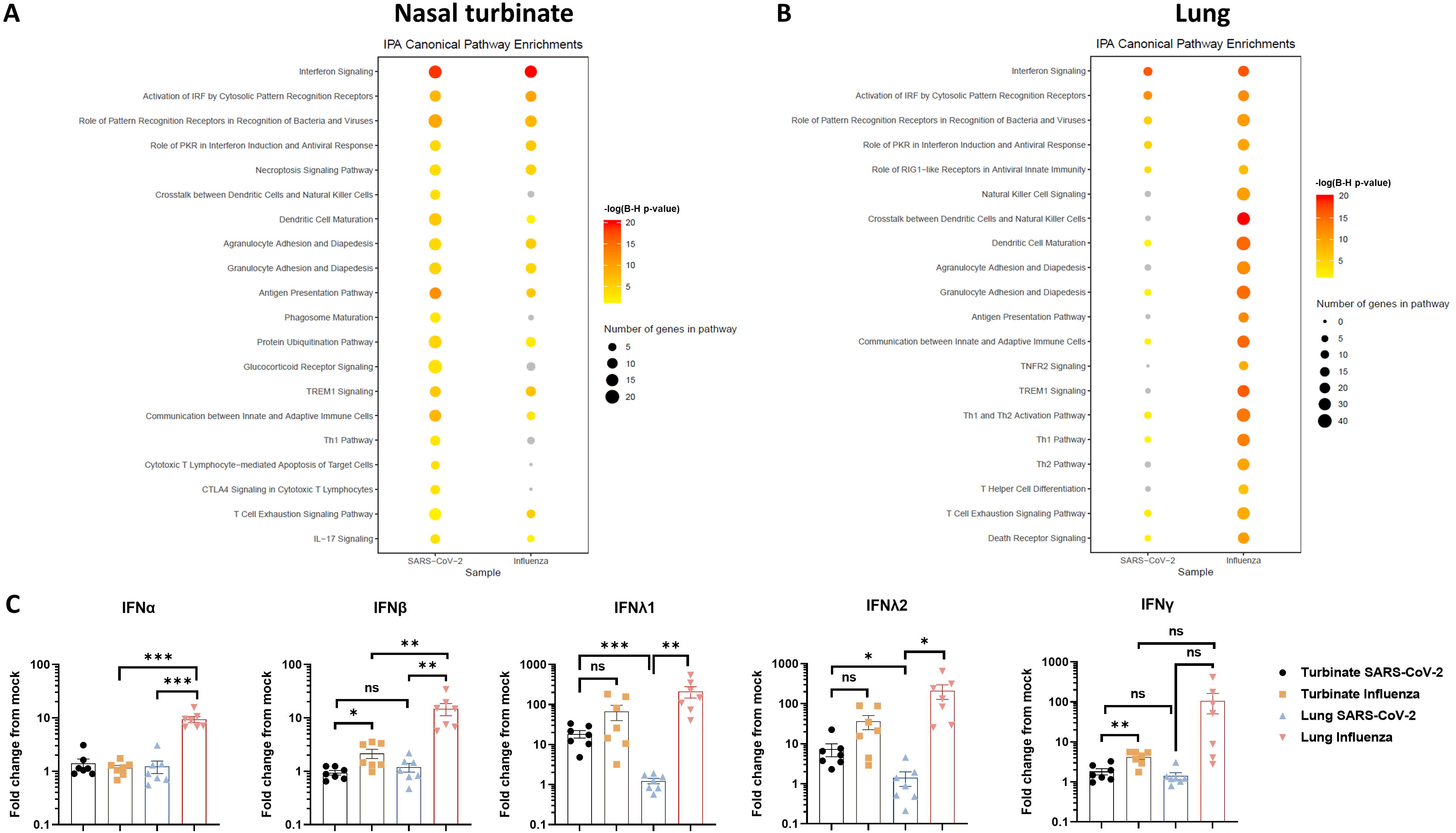
SARS-CoV-2- and influenza virus-modulated canonical pathways and interferon gene expression in nasal turbinate versus lung tissues. (**A,B**) Dot plots of selected IPA® canonical pathways in nasal turbinate (**A**) and lung tissues (**B**). The size of the dot corresponds to the number of the significantly differentially expressed genes that participate in the pathway, and the colour is according to –log(B-H P value). (**C**) Interferon responses of nasal turbinate and lung tissues, infected with SARS-CoV-2 or Influenza virus. RNA from infected and mock-infected tissues was extracted at 24 hours post infection, and analysed for the indicated interferon mRNA expression by qRT-PCR, normalized by the house keeping gene β actin. The results are presented as fold-change from mock. The results shown are representative of 7 independent nasal turbinate tissues and 7 independent lung tissues from different individuals. Significant differences are indicated by *(*P* < .05), **(*P* < .01), or ***(*P* < .001); ns, non-significant.

Influenza virus infection of the same nasal turbinate tissues differentially affected the expression of a lower number of genes compared to SARS-CoV-2 (186; 182 upregulated and 4 downregulated; Figure 3B). Comparison between the nasal turbinate tissue response to SARS-CoV-2 and influenza, identified 173 common and 198 SARS-CoV-2-distinct differentially expressed (DE) genes (Figure 3B). Innate immunity genes related to interferon signaling, immune activation, and antiviral pathways were commonly induced (albeit to a variable extent) by the 2 viruses in the nasal mucosal tissues (see also Figure 5A). The common and distinct response patterns of the nasal mucosal tissues to SARS-CoV-2 and influenza virus could be clearly delineated by a clustered heatmap analysis of all significantly DE genes, which identified 4 clusters of genes, defined by the direction (upregulation vs. downregulation) and/or the extent of their differential expression (Figure 3D). In general, the nasal turbinate tissue response to SARS-CoV-2 appeared broader than the response to influenza virus, and included innate immune and immune cell maturation and activation pathways which were distinctively or more significantly enriched following SARS-CoV-2 infection (Figures 3D, 5A). We also noted the more significant effect of SARS-CoV-2 on pathways of translation regulation (Figures 3D) – a finding which could be related to the multi-faceted strategies employed by coronaviruses in general and by SARS-CoV-2 to suppress host protein synthesis (38).

#### Innate immune responses of the infected lung tissues

Strikingly, a different response pattern was observed upon infection of the lung tissues, where SARS-CoV-2 (despite higher infection level - as demonstrated by qRT-PCR and percent viral reads; Figures 2A, S1) elicited a restricted tissue response. This was reflected in the PCA analysis, showing that influenza virus-infected tissues could be grouped by their greater transcriptional perturbation (compared to both mock- and SARS-CoV-2 infected samples) along PC1, whereas SARS-CoV-2 elicited modest transcriptional changes in this space (Figure 4A). The relatively restricted lung tissue response to SARS-CoV-2 was demonstrated by the lower number of differentially-expressed genes (124 genes; 117 upregulated and 7 downregulated; Figure 4B), along with the lower extent of gene upregulation (Figure 4C,D), compared to the effect of SARS-CoV-2 in nasal tissues (Figure 3) and to the parallel effect of influenza virus in the same lung tissues (which resulted in a response of much greater magnitude, affecting 1072 genes, the majority of which were not affected by SARS-CoV-2 infection; Figure 3B). In fact, in agreement with the PCA (Figure 4A), a clustered heatmap analysis of all significantly DE genes, showed that the relative transcriptional profile of SARS-CoV-2 infected lung tissues was closer to that of mock-infected tissues, and completely distinct from that of the highly responsive influenza-infected lung tissues (Figure 4D). While both viruses induced the expression of genes related to interferon signaling pathway (Figures 4E, 5B), influenza virus infection of the lungs induced a wide range of innate immune, immune cell activation differentiation and trafficking signaling pathways (Figures 4F,G, 5B), cytokines and chemokines (Figures S2, S3), not affected by SARS-CoV-2.

Employing independent qRT-PCR, we showed a low-to-absent upregulation of IFN-I, IFN-II, and IFN-III by SARS-CoV-2, compared to influenza virus, with the differences reaching statistical significance for IFNα, IFNβ and IFNλ (Figure 5C). Significantly, no upregulation of IFNλ1 was observed in all SARS-CoV-2 infected lung tissues examined, a finding that contrasted with the upregulation of IFNλ1 in both SARS-CoV-2-infected turbinate tissues and influenza-infected lung tissues (Figure 5C).

Together, the combined data revealed that SARS-CoV-2 affected the nasal mucosal tissues and the lung tissues in a distinct virus-specific and organ-specific manner. SARS-CoV-2 triggered a robust antiviral and proinflammatory innate immune response, broader than the innate response to influenza virus infection, in the nasal turbinate tissues, yet induced a restricted innate immune response and no apparent IFN induction in the lung tissues, which was further underscored by the vigorous response of the same lung tissues to influenza virus.

## DISCUSSION

We have characterized the initial steps of SARS-CoV-2 infection and innate immune response within the natural complexity of human nasal turbinate tissues, maintained in organ culture. Comparing the global transcriptional signatures of nasal and lung tissues, infected in parallel with SARS-CoV-2 and influenza virus, we have revealed for the first time distinct virus-host interactions in the upper and lower respiratory tract, which could determine the outcome and pathogenesis of COVID-19. SARS-CoV-2 productively infected the nasal turbinate tissues (Figures 1,2); Consistent with the expression pattern of the viral receptors ACE2 and TMPRSS2, we showed the tropism of SARS-CoV-2 to respiratory epithelial cells lining the nasal mucosa (Figure 1). Active viral replication was demonstrated by the rapid increase in viral sub-genomic mRNA within the infected tissues, along with de-novo secretion of infectious viral progeny (Figure 2). Hence, the nasal mucosa, supporting efficient viral replication, could constitute a key site of transmission, from which the progeny virus may further spread to the lungs (across the respiratory mucosa, or more likely via aspiration from the infected nasal secretions), and to the central nervous system [via the olfactory neural–mucosal interface; (39)].

Importantly, employing gene-wide transcriptome analysis, we have shown that the nasal mucosal tissue mounted a robust antiviral and inflammatory innate immune response to SARS-CoV-2, with upregulation of numerous antiviral ISGs, cytokines, and chemokines, related to interferon signaling and immune cell activation pathways (Figures 3,5,S2,S3). Moreover, comparative analysis of the nasal tissue innate response to SARS-CoV-2 and influenza virus, while demonstrating commonly induced antiviral pathways, identified virus-specific transcriptional footprints, with SARS-CoV-2 inducing an overall broader nasal-mucosal innate responses than influenza virus (Figures 3,5,S2).

In sharp contrast, infected lung tissues exhibited a restricted response to SARS-CoV-2 infection, despite comparable-to-higher viral infection levels. This finding was further underscored by the strong innate immune response of the same tissues to influenza virus infection. While the interferon signaling pathway was commonly induced by the two viruses, a wide range of antiviral and immune-cell activation pathways, cytokines and chemokines, which were induced following influenza virus infection were not stimulated by SARS-CoV-2 infection in the lungs (Figures 4, 5, S2, S3). The restricted lung-tissue response observed herein is in agreement with and expands recent reports of silenced IFN response to SARS-CoV-2 in transformed and primary human airway epithelial cell cultures and in lung explants, whereas some reports demonstrated specific chemokines induction in primary cells or even IFN elevations in lung organoids (9–12, 25, 26). These differences reflect the complex interplay between the virus and the different cell types under varying experimental conditions in culture and within integral multi-cell-type tissues.

Our findings imply that SARS-CoV-2 successfully manipulates the innate immune response in the lung tissues, which were otherwise capable of mounting a robust IFN antiviral response to influenza. In this regard, SARS-CoV-2 has been shown to encode synergistic innate immune antagonist genes [i.e., Nsp1-shutting down cellular translation, Nsp3, Nsp5, Nsp10, Nsp13, Nsp14, ORF3, ORF6 and ORF7, ORF8; (12, 38, 40, 41)], and thus may more effectively dampen the lung antiviral defence compared with influenza virus, whose IFN evasion function is mediated mainly by NS1 (12, 42). The question remains: why the same SARS-CoV-2 immune manipulation strategies are rendered ineffective in the nasal mucosa tissue milieu?

We suggest that the nasal mucosa, being constantly exposed to environmental agents and resident microflora (unlike the relatively sterile lower respiratory tract), is conditioned to persistent innate immune signalling, which could override the viral antagonists. In support of this hypothesis is the recently demonstrated skewed expression of innate immune genes in cultured nasal epithelial cells (13).

A schematic illustration of the differential tissue-specific innate immune responses to SARS-CoV-2 in nasal and lung tissues, as compared to influenza virus mediated responses, is shown in Figure 6. Our findings highlight the potential importance of the nasal mucosa as a first-barrier to SARS-CoV-2 infection. However, once the virus gains access to the lungs, the compromised early innate immune response could impede viral clearance. Beyond the global transcriptomics pattern, the conspicuous lack of type I and III IFNs upregulation (with an absolute lack of INFλ1) in early SARS-CoV-2-infected lung tissues (as opposed to their significant stimulation in influenza-infected lung tissues; Figures 5C) deserves further consideration, in relation to the distinctive late-phase pathogenesis of COVID-19; IFNs type I and III share common antiviral functions, yet, type III IFNs, beyond their role in antiviral defense, have been shown to exert critical immune-regulatory activities -limiting excessive local inflammation (15). Thus, it is tempting to speculate that the restricted antiviral and immune-regulatory IFN response in early SARS-CoV-2-infected lung tissues (not observed following influenza virus infection), could mechanistically explain the subsequent uncontrolled SARS-CoV-2 replication and imbalanced hyper-inflammatory response, characteristic of late-phase COVID-19. Our study has several limitations. The cellular heterogeneity within the tissues limits the resolution of isolated molecular pathways. Additionally, native respiratory tissues in organ culture are relatively short-lived (∼a week) compared to primary human airway epithelial cells and to the self-renewable stem cell-derived organoid cultures, which have proven useful for the studies of SARS-CoV-2 infection and cellular response (9–11, 43, 44). Nonetheless, our studies recapitulate viral infection and host response within the authentic multicellular and morphologically-intact tissue microenvironment - containing tissue epithelial, vascular endothelial, stromal and immune cells, and the specific extracellular matrix. It is notable that the comparative data between the human nasal and lung tissues were not obtained from the same individuals. Yet, we believe that the findings, based on extensive analyses of independent tissues from different individuals, faithfully support the generalizability of the observed tissue-specific patterns. Notwithstanding, the comparison of SARS-CoV-2 and Influenza infections was done in parallel within the same donor tissue. Furthermore, these studies capture the inherent person-to-person variability of innate immune responses, thereby paving the way to future studies of personal host features which determine the innate responses to viral infection along the respiratory tract.

**Figure 6.**
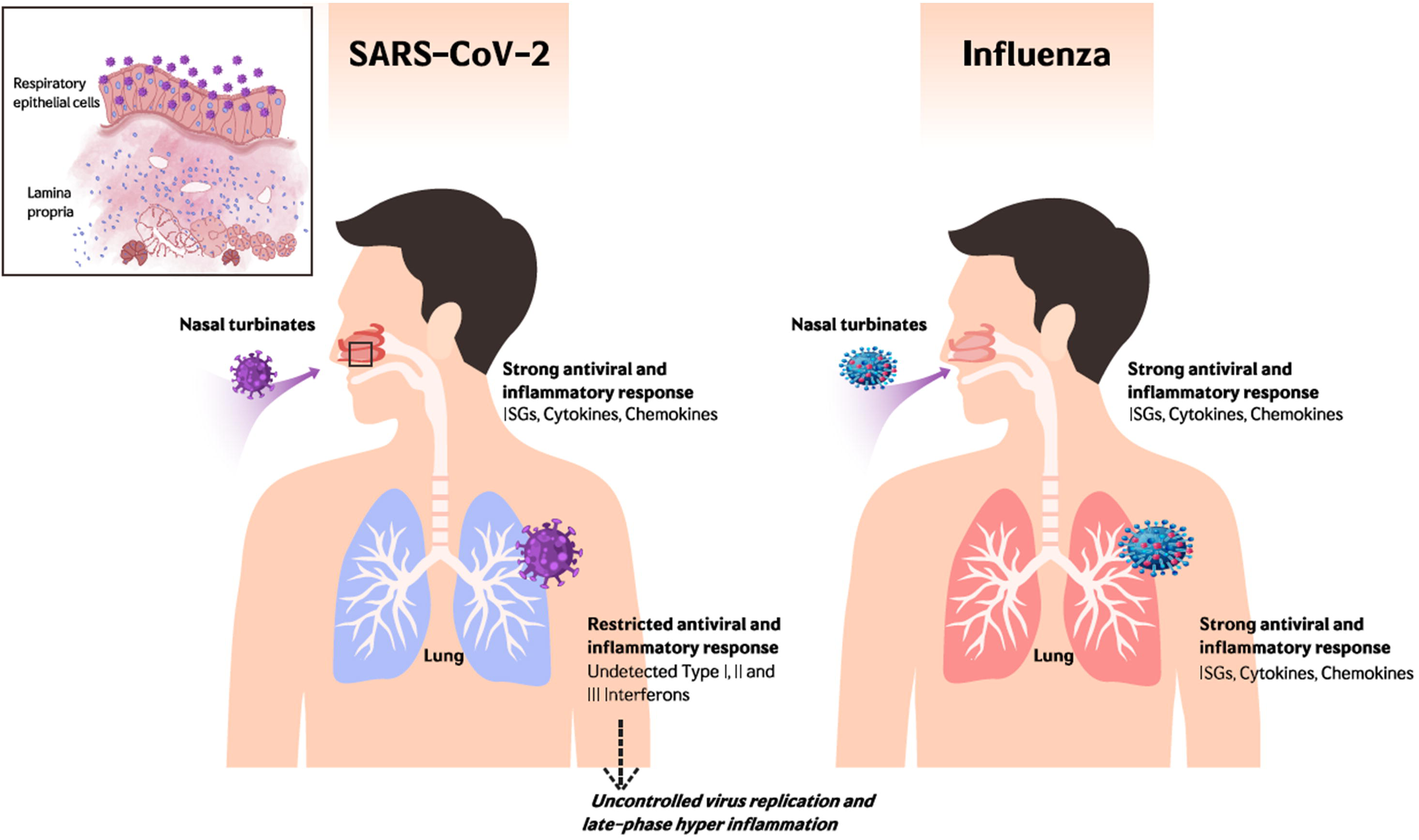
Schematic illustration of the early patterns of viral infection and local-mucosal innate immune responses in human nasal turbinate and lung tissues. In the work presented here, we show that SARS-CoV-2 productively infects respiratory epithelial cells within the nasal turbinate tissues. Comparing the innate response patterns of nasal and lung tissues, infected in parallel with SARS-CoV-2 and influenza virus, we have revealed differential tissue-specific and virus-specific innate immune responses in the upper and lower respiratory tract. Our findings emphasize the role of the nasal mucosa in viral transmission and innate antiviral defence, whereas the restricted innate immune response in early-SARS-CoV-2-infected lung tissues (contrasting with their robust response to influenza virus) could underlie the unique late-phase lung damage of advanced COVID-19. ; ISGs, Interferon stimulated genes.

In summary, we have demonstrated the active replication of SARS-CoV-2 in native human nasal-mucosal tissues, providing new insights into differential tissue-specific innate immune responses to SARS-CoV-2 in nasal and lung tissues. Our findings shed light on the nasal mucosa as a key site of viral transmission and innate immune defense, implying a window of opportunity for early interventions, whereas the restricted innate immune response in early-SARS-CoV-2-infected lung tissues, contrasting with their robust response to influenza virus, could underlie the unique extensive late-phase lung damage of advanced COVID-19. Studies in the nasal mucosal model can be further employed to assess the impact of viral evolutionary changes, and evaluate new therapeutic and preventive measures against SARS-CoV-2 and other human respiratory pathogens.

## MATERIALS AND METHODS

### Cells and viruses

Simian kidney Vero E6 (ATCC CRL-1586) and Madin-Darby Canine Kidney (MDCK, ATCC® CCL-34™) cells were maintained in Eagle’s Minimum Essential Medium (EMEM; Biological Industries, Beit Haemek, Israel), supplemented with 10% fetal bovine serum, 2 mM L-Glutamine, 10 IU/ml Penicillin, and 10 μg/ml streptomycin (Biological Industries, Beit Haemek, Israel). SARS-CoV-2 isolate USA-WA1/2020 (NR-52281; obtained from BEI resources) was propagated in Vero E6 cells. SARS-CoV-2 clinical isolate (SARS-CoV-2 isolate Israel-Jerusalem-854/2020) was isolated on Vero E6 cells from a positive nasopharyngeal swab sample, obtained at the Hadassah Hospital Clinical Virology Laboratory. The virus was isolated and propagated (3 passages) in Vero E6 cells, and sequence verified. Influenza virus A(H1N1) pdm09 (NIBRG-121xp, Cat# 09/268; obtained from NIBSC, UK) was propagated in MDCK cells. The virus titers of cleared infected cells- and tissue supernatants were determined by a standard TCID_50_ assay on Vero E6 cells (SARS-CoV-2) or MDCK cells (influenza virus).

### Preparation and infection of nasal turbinate and lung organ cultures

Nasal turbinate and lung organ cultures were prepared and infected as previously described (27, 36). In brief, inferior nasal turbinate tissues were obtained from consented patients undergoing turbinectomy procedures, and lung tissues (the tumor free margins) were obtained from consented patients undergoing lobectomy operations. The studies were approved by the Hadassah Medical Center and the Sheba Medical Center Institutional Review Boards. Fresh tissues were kept on ice until further processed at the same day. The tissues were sectioned by a microtome (McIlwain Tissue Chopper; Ted Pella, INC.) into thin slices (250 μ -thick slices; each encompassing ∼10 cell layers), and incubated in 0.3 ml of enriched RPMI medium (for the nasal turbinate tissues) or DMEM/F-12 medium with MEM Vitamin Solution (for the lung tissues), with 10% fetal bovine serum, 2.5 μg glucose/ml, 2 mM glutamine, 10 IU/ml penicillin, 10 μg/ml streptomycin, and 0.25 μ ml amphotericin B, at 37°C, 5% CO2. The tissues were processed and infected at the same day (the day of harvesting; Day 0). For infection of the organ cultures, the tissues were placed in 48-well plates and inoculated with the respective virus (2×10^5^ TCID_50_/well in 0.3 ml) for 12h to allow effective viral adsorption. Following viral adsorption, the cultures were washed three times (in 0.3 ml of complete medium) and further incubated for the duration of the experiment, with replacement of the culture medium every 2 to 3 days. Tissue viability was monitored by the mitochondrial dehydrogenase enzyme (MTT) assay as previously described (28). All infection and tissue processing experiments were performed in a BSL-3 facility.

### Whole-mount tissue immunofluorescence

Tissues were fixed in Image-iT™ Fixative Solution (4% formaldehyde, methanol-free; Thermo Fisher Scientific, Cat# R37814) for hours, washed in PBS and transferred to 80% ethanol. The tissues were permeabilized by 0.3% Triton-X100 in PBS (PBST) and further incubated with Animal-Free Blocker® (Vector laboratories, Cat# SP-5035-100) to block nonspecific antibody binding, followed by incubation with the primary antibodies in Animal-Free Blocker® at room temperature overnight. The tissues were than washed 4 times in PBST, incubated with the secondary antibodies in Animal-Free Blocker® for at room temperature overnight, washed 4 times with PBST, and incubated with 4’,6-diamidino-2-phenylindole (DAPI, 10uM, Abcam, Cat# ab228549) as a nuclear stain. The following primary antibodies were used: Ep-CAM (Mouse monoclonal, 1:100, Thermo Fisher Scientific, Cat# 14-9326-82; for the detection of epithelial cells), ACE2 (Rabbit monoclonal, 1:100, Thermo Fisher Scientific, Cat# MA5-32307), TMPRSS2 (Rabbit Polyclonal, 1:50, Sino biological, Cat# 204314-T08), SARS-CoV-2 Nucleocapsid (Rabbit Polyclonal, 1:50, Novus Biologicals, Cat# NB100-56576), and Influenza A Nucleoprotein (Goat polyclonal, 1:100, Abcam, Cat# ab20841). The following secondary antibodies were used: Donkey anti-Mouse IgG pre-adsorbed, Alexa Fluor® 568 (1:250, Abcam, Cat# ab175700), Donkey anti-Goat IgG pre-adsorbed, Alexa Fluor® 647 (1:250, Abcam, Cat# ab150135), Goat anti-Rabbit IgG Highly Cross-Adsorbed Alexa Fluor Plus 647 (1:250, Thermo Fisher Scientific, Cat# A32733). For tissue clearing, stained preparations were dehydrated with 100% Ethanol for 1h, and later submerged and mounted in ethyl-cinnamate (99%; Sigma, Cat# 112372) as previously described (45). Whole-mount tissues were visualized using a Nikon A1R confocal microscope and were analyzed using NIS Elements software (Nikon).

### RNA purification and quantification

Infected- and mock-infected organ cultures and the respective supernatants were flash-frozen and stored at -80°C until assayed. RNA was extracted using NucleoSpin RNA Mini kit for RNA purification (Macherey-Nagel, Cat #740955.250) according to the manufacturer’s instructions, and subjected to reverse transcription, using High-Capacity cDNA Reverse Transcription Kit (Thermo Fisher Scientific, Cat#). Quantitative real time (RT)-PCR was performed on a Quantstudio 3™ (Thermo Fisher Scientific) instrument, using Fast SYBR™ Green Master Mix (Thermo Fisher Scientific, Cat# 4385614), or TaqMan™ Fast Advanced Master Mix (Thermo Fisher Scientific, Cat# 4444558). The employed primers and probe sequences are listed in Table S1.

### cDNA library preparation, deep sequencing, and bioinformatics analysis

Nasal organ cultures (from 3 independent donors) and lung organ cultures (from 5 independent donors) were each infected in parallel with SARS-CoV-2 and influenza virus (with mock-infected controls used for each individual tissue).

Preparation of transcriptome libraries and deep sequencing were performed as previously described in detail (35). Normalization and differential expression were done with the DESeq2 package (version 1.22.2). Samples of each tissue were analyzed separately, after removing genes with less than 10 counts. Differential expression was calculated with default parameters, incorporating both the donor and the infection information in the statistical model. Pairwise comparisons of the infected-versus mock-infected samples were performed, while considering the donor effect (design = ∼ Donor + Infection). Genes with a significant effect (padj<0.1) were further filtered based on their baseMean and log2FoldChange values, requiring baseMean > 5 and |log2FoldChange| > 5/baseMean^0.5 + 0.3. In order test the contribution of infection to the variance in the system as a whole, the LRT test was used, comparing the full model, Donor + Infection, to a reduced model of just the Donor factor. Genes with a padj<0.05 in this test were taken as significant and clustered using R’s kmeans function. Results are shown for k=2 in lungs tissue and for k=4 in turbinates tissue.

Pathway and molecular function and disease enrichment analysis of the significantly differentially expressed genes was carried out using the Ingenuity Pathway Analysis (IPA®) (QIAGEN Inc., https://digitalinsights.qiagen.com/products-overview/discovery-insights-portfolio/content-exploration-and-databases/qiagen-ipa/).

Dot plots of selected IPA® canonical pathways (based on IPA® values for BH P-values and number of genes) were generated using ggplot2 R graphical package (Wickham H (2016). ggplot2: Elegant Graphics for Data Analysis. Springer-Verlag New York. ISBN 978-3-319-24277-4, https://ggplot2.tidyverse.org.)

### Statistical analysis

All data, presented as means ± standard errors of the mean (SEM), were analyzed using unpaired, two-tailed Student’s t test in GraphPad Prism 8 software (GraphPad Software Inc., San Diego CA). P values of <0.05 were considered significant. Statistical analysis of the transcriptome data was done as described above.

### Data availability

The transcriptomic data described in this publication have been deposited in the NCBI Gene Expression Omnibus and are accessible through GEO series accession number GSE163959.

## ACKNOWLEGEMENTS

This work was supported by the Israel Science Foundation [530/18], The Israeli Science Ministry, The Rothschild Foundation, the European Union Seventh Framework Program ERA-NET Infect-ERA CYMAF consortium, and by the Samueli Foundation.

We thank Dr. Hadar Benyamini and Adi Turjeman for their help in the transcriptome analysis. We thank Dr. Adi Stern and Noam Harel for sequencing the SARS-CoV-2 clinical isolate.

The authors declare no conflict of interest.

## SUPPELEMENTAL MATERIAL

**Supplemental Figure S1. Percent of viral transcripts in the transcriptome of infected tissues.** RNA was extracted from SARS-CoV-2 or influenza A(H1N1) pdm09 virus-infected turbinate and lung tissues at 24 hours post infection and subjected to transcriptome analysis. Three independent donors’ turbinate tissues and five independent donors’ lung tissues (with two biological replicates for each experimental condition) were included in the analysis. Mean percent values of viral transcripts (out of all the sequence reads in the transcriptome) with SEM are shown. ** denotes P<0.01.

**Supplemental Figure S2. Heatmaps representing cytokine and chemokine expression levels**. Nasal turbinate (**A**) and lung (**B**) organ cultures were mock-infected or infected in parallel with SARS-CoV-2 or influenza A(H1N1) pdm09 (2×10^5^ TCID_50_/well). At 24 hours post infection, RNA was extracted and subjected to transcriptome analysis. Three independent nasal turbinate donors’ tissues and five independent lung donors’ tissues (with two biological replicates for each experimental condition) were included in the analysis. The expression of the cytokines and chemokines with the lowest p-value (for SARS-CoV-2-infected nasal turbinate tissues and for influenza virus-infected lung tissues) is shown. Normalized expression values were scaled at gene level, then hierarchically clustered and drawn as a heatmap. The scale is shown at top-right and the clustering order is given on the left.

**Supplemental Figure S3. Effect of SARS-CoV-2 and influenza virus infection on the expression of selected innate immunity genes in nasal turbinate and lung organ cultures.** RNA from SARS-CoV-2-, influenza A(H1N1) pdm09-, and mock-infected cultures was extracted at 24 hours post infection and analysed for the indicated gene expression by qRT-PCR, normalized by the expression of the house keeping gene β actin.

## REFERENCES

1. Huang C, Wang Y, Li X, Ren L, Zhao J, Hu Y, Zhang L, Fan G, Xu J, Gu X, Cheng Z, Yu T, Xia J, Wei Y, Wu W, Xie X, Yin W, Li H, Liu M, Xiao Y, Gao H, Guo L, Xie J, Wang G, Jiang R, Gao Z, Jin Q, Wang J, Cao B. 2020. Clinical features of patients infected with 2019 novel coronavirus in Wuhan, China. Lancet 395:497–506.

2. Bradley BT, Maioli H, Johnston R, Chaudhry I, Fink SL, Xu H, Najafian B, Deutsch G, Lacy JM, Williams T, Yarid N, Marshall DA. 2020. Histopathology and ultrastructural findings of fatal COVID-19 infections in Washington State: a case series. Lancet 396:320–332.

3. Vardhana SA, Wolchok JD. 2020. The many faces of the anti-COVID immune response. J Exp Med 217:1–10.

4. Wauters E, Thevissen K, Wouters C, Bosisio FM, De Smet F, Gunst J, Humblet-Baron S, Lambrechts D, Liston A, Matthys P, Neyts J, Proost P, Weynand B, Wauters J, Tejpar S, Garg AD. 2020. Establishing a Unified COVID-19 “Immunome”: Integrating Coronavirus Pathogenesis and Host Immunopathology. Front Immunol 11:1–5.

5. Xu Z, Shi L, Wang Y, Zhang J, Huang L, Zhang C, Liu S, Zhao P, Liu H, Zhu L, Tai Y, Bai C, Gao T, Song J, Xia P, Dong J, Zhao J, Wang FS. 2020. Pathological findings of COVID-19 associated with acute respiratory distress syndrome. Lancet Respir Med 8:420–422.

6. Hoffmann M, Kleine-Weber H, Schroeder S, Krüger N, Herrler T, Erichsen S, Schiergens TS, Herrler G, Wu NH, Nitsche A, Müller MA, Drosten C, Pöhlmann S. 2020. SARS-CoV-2 Cell Entry Depends on ACE2 and TMPRSS2 and Is Blocked by a Clinically Proven Protease Inhibitor. Cell 181:271–280.e8.

7. Hou YJ, Okuda K, Edwards CE, Martinez DR, Asakura T, Dinnon KH, Kato T, Lee RE, Yount BL, Mascenik TM, Chen G, Olivier KN, Ghio A, Tse L V., Leist SR, Gralinski LE, Schäfer A, Dang H, Gilmore R, Nakano S, Sun L, Fulcher ML, Livraghi-Butrico A, Nicely NI, Cameron M, Cameron C, Kelvin DJ, de Silva A, Margolis DM, Markmann A, Bartelt L, Zumwalt R, Martinez FJ, Salvatore SP, Borczuk A, Tata PR, Sontake V, Kimple A, Jaspers I, O’Neal WK, Randell SH, Boucher RC, Baric RS. 2020. SARS-CoV-2 Reverse Genetics Reveals a Variable Infection Gradient in the Respiratory Tract. Cell 182:429–446.e14.

8. Ziegler CGK, Allon SJ, Nyquist SK, Mbano IM, Miao VN, Tzouanas CN, Cao Y, Yousif AS, Bals J, Hauser BM, Feldman J, Muus C, Wadsworth MH, Kazer SW, Hughes TK, Doran B, Gatter GJ, Vukovic M, Taliaferro F, Mead BE, Guo Z, Wang JP, Gras D, Plaisant M, Ansari M, Angelidis I, Adler H, Sucre JMS, Taylor CJ, Lin B, Waghray A, Mitsialis V, Dwyer DF, Buchheit KM, Boyce JA, Barrett NA, Laidlaw TM, Carroll SL, Colonna L, Tkachev V, Peterson CW, Yu A, Zheng HB, Gideon HP, Winchell CG, Lin PL, Bingle CD, Snapper SB, Kropski JA, Theis FJ, Schiller HB, Zaragosi LE, Barbry P, Leslie A, Kiem HP, Flynn JAL, Fortune SM, Berger B, Finberg RW, Kean LS, Garber M, Schmidt AG, Lingwood D, Shalek AK, Ordovas-Montanes J, Banovich N, Brazma A, Desai T, Duong TE, Eickelberg O, Falk C, Farzan M, Glass I, Haniffa M, Horvath P, Hung D, Kaminski N, Krasnow M, Kuhnemund M, Lafyatis R, Lee H, Leroy S, Linnarson S, Lundeberg J, Meyer K, Misharin A, Nawijn M, Nikolic MZ, Pe’er D, Powell J, Quake S, Rajagopal J, Tata PR, Rawlins EL, Regev A, Reyfman PA, Rojas M, Rosen O, Saeb-Parsy K, Samakovlis C, Schiller H, Schultze JL, Seibold MA, Shepherd D, Spence J, Spira A, Sun X, Teichmann S, Theis F, Tsankov A, van den Berge M, von Papen M, Whitsett J, Xavier R, Xu Y, Zhang K. 2020. SARS-CoV-2 Receptor ACE2 Is an Interferon-Stimulated Gene in Human Airway Epithelial Cells and Is Detected in Specific Cell Subsets across Tissues. Cell 181:1016–1035.e19.

9. Vanderheiden A, Ralfs P, Chirkova T, Upadhyay AA, Zimmerman MG, Bedoya S, Aoued H, Tharp GM, Pellegrini KL, Manfredi C, Sorscher E, Mainou B, Lobby JL, Kohlmeier JE, Lowen AC, Shi P-Y, Menachery VD, Anderson LJ, Grakoui A, Bosinger SE, Suthar MS. 2020. Type I and Type III Interferons Restrict SARS-CoV-2 Infection of Human Airway Epithelial Cultures. J Virol 94.

10. Fiege JK, Thiede JM, Nanda H, Matchett WE, Moore PJ, Montanari NR, Thielen BK, Daniel J, Stanley E, Hunter RC, Menachery VD, Shen SS, Bold TD, Langlois RA. 2020. Single cell resolution of SARS-CoV-2 tropism, antiviral responses, and susceptibility to therapies in primary human airway epithelium. bioRxiv 2020.10.19.343954.

11. Katsura H, Sontake V, Tata A, Kobayashi Y, Edwards CE, Heaton BE, Konkimalla A, Asakura T, Mikami Y, Fritch EJ, Lee PJ, Heaton NS, Boucher RC, Randell SH, Baric RS, Tata PR. 2020. Human Lung Stem Cell-Based Alveolospheres Provide Insights into SARS-CoV-2-Mediated Interferon Responses and Pneumocyte Dysfunction. Cell Stem Cell 1–15.

12. Blanco-Melo D, Nilsson-Payant BE, Liu WC, Uhl S, Hoagland D, Møller R, Jordan TX, Oishi K, Panis M, Sachs D, Wang TT, Schwartz RE, Lim JK, Albrecht RA, TenOever BR. 2020. Imbalanced Host Response to SARS-CoV-2 Drives Development of COVID-19. Cell 181:1036–1045.e9.

13. Sungnak W, Huang N, Bécavin C, Berg M, Queen R, Litvinukova M, Talavera-López C, Maatz H, Reichart D, Sampaziotis F, Worlock KB, Yoshida M, Barnes JL, Banovich NE, Barbry P, Brazma A, Collin J, Desai TJ, Duong TE, Eickelberg O, Falk C, Farzan M, Glass I, Gupta RK, Haniffa M, Horvath P, Hubner N, Hung D, Kaminski N, Krasnow M, Kropski JA, Kuhnemund M, Lako M, Lee H, Leroy S, Linnarson S, Lundeberg J, Meyer KB, Miao Z, Misharin A V., Nawijn MC, Nikolic MZ, Noseda M, Ordovas-Montanes J, Oudit GY, Pe’er D, Powell J, Quake S, Rajagopal J, Tata PR, Rawlins EL, Regev A, Reyfman PA, Rozenblatt-Rosen O, Saeb-Parsy K, Samakovlis C, Schiller HB, Schultze JL, Seibold MA, Seidman CE, Seidman JG, Shalek AK, Shepherd D, Spence J, Spira A, Sun X, Teichmann SA, Theis FJ, Tsankov AM, Vallier L, van den Berge M, Whitsett J, Xavier R, Xu Y, Zaragosi LE, Zerti D, Zhang H, Zhang K, Rojas M, Figueiredo F. 2020. SARS-CoV-2 entry factors are highly expressed in nasal epithelial cells together with innate immune genes. Nat Med 26:681– 687.

14. MacMicking JD. 2012. Interferon-inducible effector mechanisms in cell-autonomous immunity. Nat Rev Immunol 12:367–382.

15. Galani IE, Triantafyllia V, Eleminiadou EE, Koltsida O, Stavropoulos A, Manioudaki M, Thanos D, Doyle SE, Kotenko S V., Thanopoulou K, Andreakos E. 2017. Interferon-λ Mediates Non-redundant Front-Line Antiviral Protection against Influenza Virus Infection without Compromising Host Fitness. Immunity 46:875–890.e6.

16. Van Der Made CI, Simons A, Schuurs-Hoeijmakers J, Van Den Heuvel G, Mantere T, Kersten S, Van Deuren RC, Steehouwer M, Van Reijmersdal S V., Jaeger M, Hofste T, Astuti G, Corominas Galbany J, Van Der Schoot V, Van Der Hoeven H, Hagmolen Of Ten Have W, Klijn E, Van Den Meer C, Fiddelaers J, De Mast Q, Bleeker-Rovers CP, Joosten LAB, Yntema HG, Gilissen C, Nelen M, Van Der Meer JWM, Brunner HG, Netea MG, Van De Veerdonk FL, Hoischen A. 2020. Presence of Genetic Variants among Young Men with Severe COVID-19. JAMA - J Am Med Assoc 324:663–673.

17. Zhang Q, Bastard P, Liu Z, Le Pen J, Moncada-Velez M, Chen J, Ogishi M, Sabli IKD, Hodeib S, Korol C, Rosain J, Bilguvar K, Ye J, Bolze A, Bigio B, Yang R, Arias AA, Zhou Q, Zhang Y, Onodi F, Korniotis S, Karpf L, Philippot Q, Chbihi M, Bonnet-Madin L, Dorgham K, Smith N, Schneider WM, Razooky BS, Hoffmann HH, Michailidis E, Moens L, Han JE, Lorenzo L, Bizien L, Meade P, Neehus AL, Ugurbil AC, Corneau A, Kerner G, Zhang P, Rapaport F, Seeleuthner Y, Manry J, Masson C, Schmitt Y, Schlüter A, Le Voyer T, Khan T, Li J, Fellay J, Roussel L, Shahrooei M, Alosaimi MF, Mansouri D, Al-Saud H, Al-Mulla F, Almourfi F, Al-Muhsen SZ, Alsohime F, Turki S Al, Hasanato R, Van De Beek D, Biondi A, Bettini LR, D’Angio’ M, Bonfanti P, Imberti L, Sottini A, Paghera S, Quiros-Roldan E, Rossi C, Oler AJ, Tompkins MF, Alba C, Vandernoot I, Goffard JC, Smits G, Migeotte I, Haerynck F, Soler-Palacin P, Martin-Nalda A, Colobran R, Morange PE, Keles S, Çölkesen F, Ozcelik T, Yasar KK, Senoglu S, Karabela ŞN, Rodríguez-Gallego C, Novelli G, Hraiech S, Tandjaoui-Lambiotte Y, Duval X, Laouénan C, Snow AL, Dalgard CL, Milner JD, Vinh DC, Mogensen TH, Marr N, Spaan AN, Boisson B, Boisson-Dupuis S, Bustamante J, Puel A, Ciancanelli MJ, Meyts I, Maniatis T, Soumelis V, Amara A, Nussenzweig M, García-Sastre A, Krammer F, Pujol A, Duffy D, Lifton RP, Zhang SY, Gorochov G, Béziat V, Jouanguy E, Sancho-Shimizu V, Rice CM, Abel L, Notarangelo LD, Cobat A, Su HC, Casanova JL. 2020. Inborn errors of type I IFN immunity in patients with life-threatening COVID-19. Science (80-) 370:1–9.

18. Bastard P, Rosen LB, Zhang Q, Zhang Y, Dorgham K, Béziat V, Puel A, Lorenzo L, Bizien L, Assant S, Fillipot Q, Seeleuthner Y, Hadjadj J, Bigio B, Michael S, Shaw E, Chauvin SD, Belot A, Rieux-laucat F. 2020. Autoantibodies against type I IFNs in patients with. Science (80-) 4585:1–19.

19. Hadjadj J, Yatim N, Barnabei L, Corneau A, Boussier J, Smith N, Péré H, Charbit B, Bondet V, Chenevier-Gobeaux C, Breillat P, Carlier N, Gauzit R, Morbieu C, Pène F, Marin N, Roche N, Szwebel TA, Merkling SH, Treluyer JM, Veyer D, Mouthon L, Blanc C, Tharaux PL, Rozenberg F, Fischer A, Duffy D, Rieux-Laucat F, Kernéis S, Terrier B. 2020. Impaired type I interferon activity and inflammatory responses in severe COVID-19 patients. Science (80-) 369:718–724.

20. Mehta P, McAuley DF, Brown M, Sanchez E, Tattersall RS, Manson JJ. 2020. COVID-19: consider cytokine storm syndromes and immunosuppression. Lancet 395:1033–1034.

21. Acharya D, Liu GQ, Gack MU. 2020. Dysregulation of type I interferon responses in COVID-19. Nat Rev Immunol 20:397–398.

22. Muñoz-Fontela C, Dowling WE, Funnell SGP, Gsell PS, Riveros-Balta AX, Albrecht RA, Andersen H, Baric RS, Carroll MW, Cavaleri M, Qin C, Crozier I, Dallmeier K, de Waal L, de Wit E, Delang L, Dohm E, Duprex WP, Falzarano D, Finch CL, Frieman MB, Graham BS, Gralinski LE, Guilfoyle K, Haagmans BL, Hamilton GA, Hartman AL, Herfst S, Kaptein SJF, Klimstra WB, Knezevic I, Krause PR, Kuhn JH, Le Grand R, Lewis MG, Liu WC, Maisonnasse P, McElroy AK, Munster V, Oreshkova N, Rasmussen AL, Rocha-Pereira J, Rockx B, Rodríguez E, Rogers TF, Salguero FJ, Schotsaert M, Stittelaar KJ, Thibaut HJ, Tseng C Te, Vergara-Alert J, Beer M, Brasel T, Chan JFW, García-Sastre A, Neyts J, Perlman S, Reed DS, Richt JA, Roy CJ, Segalés J, Vasan SS, Henao-Restrepo AM, Barouch DH. 2020. Animal models for COVID-19. Nature 586:509–515.

23. Yang L, Han Y, Nilsson-Payant BE, Gupta V, Wang P, Duan X, Tang X, Zhu J, Zhao Z, Jaffré F, Zhang T, Kim TW, Harschnitz O, Redmond D, Houghton S, Liu C, Naji A, Ciceri G, Guttikonda S, Bram Y, Nguyen DHT, Cioffi M, Chandar V, Hoagland DA, Huang Y, Xiang J, Wang H, Lyden D, Borczuk A, Chen HJ, Studer L, Pan FC, Ho DD, tenOever BR, Evans T, Schwartz RE, Chen S. 2020. A Human Pluripotent Stem Cell-based Platform to Study SARS-CoV-2 Tropism and Model Virus Infection in Human Cells and Organoids. Cell Stem Cell 27:125–136.e7.

24. Clausen TM, Sandoval DR, Spliid CB, Pihl J, Perrett HR, Painter CD, Narayanan A, Majowicz SA, Kwong EM, McVicar RN, Thacker BE, Glass CA, Yang Z, Torres JL, Golden GJ, Bartels PL, Porell RN, Garretson AF, Laubach L, Feldman J, Yin X, Pu Y, Hauser BM, Caradonna TM, Kellman BP, Martino C, Gordts PLSM, Chanda SK, Schmidt AG, Godula K, Leibel SL, Jose J, Corbett KD, Ward AB, Carlin AF, Esko JD. 2020. SARS-CoV-2 Infection Depends on Cellular Heparan Sulfate and ACE2. Cell 183:1043–1057.e15.

25. Hui KPY, Cheung MC, Perera RAPM, Ng KC, Bui CHT, Ho JCW, Ng MMT, Kuok DIT, Shih KC, Tsao SW, Poon LLM, Peiris M, Nicholls JM, Chan MCW. 2020. Tropism, replication competence, and innate immune responses of the coronavirus SARS-CoV-2 in human respiratory tract and conjunctiva: an analysis in *ex-vivo* and in-vitro cultures. Lancet Respir Med 8:687–695.

26. Chu H, Chan JFW, Wang Y, Yuen TTT, Chai Y, Hou Y, Shuai H, Yang D, Hu B, Huang X, Zhang X, Cai JP, Zhou J, Yuan S, Kok KH, To KKW, Chan IHY, Zhang AJ, Sit KY, Au WK, Yuen KY. 2020. Comparative replication and immune activation profiles of SARS-CoV-2 and SARS-CoV in human lungs: An *ex vivo* study with implications for the pathogenesis of COVID-19. Clin Infect Dis 71:1400–1409.

27. Massler A, Kolodkin-Gal D, Meir K, Khalaileh A, Falk H, Izhar U, Shufaro Y, Panet A. 2011. Infant lungs are preferentially infected by adenovirus and herpes simplex virus type 1 vectors: Role of the tissue mesenchymal cells. J Gene Med 13:101–113.

28. Weisblum Y, Panet A, Zakay-Rones Z, Haimov-Kochman R, Goldman-Wohl D, Ariel I, Falk H, Natanson-Yaron S, Goldberg MD, Gilad R, Lurain NS, Greenfield C, Yagel S, Wolf DG. 2011. Modeling of Human Cytomegalovirus Maternal-Fetal Transmission in a Novel Decidual Organ Culture. J Virol 85:13204–13213.

29. Yaacov B, Lazar I, Tayeb S, Frank S, Izhar U, Lotem M, Perlman R, Ben-Yehuda D, Zakay-Rones Z, Panet A. 2012. Extracellular matrix constituents interfere with Newcastle disease virus spread in solid tissue and diminish its potential oncolytic activity. J Gen Virol 93:1664–1672.

30. Kunicher N, Tzur T, Amar D, Chaouat M, Yaacov B, Panet A. 2011. Characterization of factors that determine lentiviral vector tropism in skin tissue using an *ex vivo* model. J Gene Med 13:209–220.

31. Weisblum Y, Panet A, Haimov-Kochman R, Wolf DG. 2014. Models of vertical cytomegalovirus (CMV) transmission and pathogenesis. Semin Immunopathol 36:615–625.

32. Tsalenchuck Y, Steiner I, Panet A. 2016. Innate defense mechanisms against HSV-1 infection in the target tissues, skin and brain. J Neurovirol 22:641–649.

33. Weisblum Y, Panet A, Zakay-Rones Z, Vitenshtein A, Haimov-Kochman R, Goldman-Wohl D, Oiknine-Djian E, Yamin R, Meir K, Amsalem H, Imbar T, Mandelboim O, Yagel S, Wolf DG. 2015. Human cytomegalovirus induces a distinct innate immune response in the maternal-fetal interface. Virology 485:289–296.

34. Weisblum Y, Oiknine-Djian E, Vorontsov OM, Haimov-Kochman R, Zakay-Rones Z, Meir K, Shveiky D, Elgavish S, Nevo Y, Roseman M, Bronstein M, Stockheim D, From I, Eisenberg I, Lewkowicz AA, Yagel S, Panet A, Wolf DG. 2017. Zika Virus Infects Early- and Midgestation Human Maternal Decidual Tissues, Inducing Distinct Innate Tissue Responses in the Maternal-Fetal Interface. J Virol 91:1–13.

35. Weisblum Y, Oiknine-Djian E, Zakay-Rones Z, Vorontsov O, Haimov-Kochman R, Nevo Y, Stockheim D, Yagel S, Panet A, Wolf DG. 2017. APOBEC3A Is Upregulated by Human Cytomegalovirus (HCMV) in the Maternal-Fetal Interface, Acting as an Innate Anti-HCMV Effector. J Virol 91:1–13.

36. Alfi O, From I, Yakirevitch A, Drendel M, Wolf M, Meir K, Zakay-Rones Z, Nevo Y, Elgavish S, Ilan O, Weisblum Y, Tayeb S, Gross M, Jonas W, Ives J, Oberbaum M, Panet A, Wolf DG. 2020. Human Nasal Turbinate Tissues in Organ Culture as a Model for Human Cytomegalovirus Infection at the Mucosal Entry Site. J Virol 94:1–12.

37. El-Diwany R, Soliman M, Sugawara S, Breitwieser F, Skaist A, Coggiano C, Sangal N, Chattergoon M, Bailey JR, Siliciano RF, Blankson JN, Ray SC, Wheelan SJ, Thomas DL, Balagopal A. 2018. CMPK2 and BCL-G are associated with type 1 interferon–induced HIV restriction in humans. Sci Adv 4:1–12.

38. Finkel Y, Gluck A, Winkler R, Nachshon A, Mizrahi O, Zuckerman B, Slobodin B, Yahalom-Ronen Y, Tamir H, Israely T, Paran N, Schwartz M, Stern-Ginossar N. 2020. SARS-CoV-2 utilizes a multipronged strategy to suppress host 2 protein synthesis Introduction. bioRxiv 2020.11.25.398578.

39. Meinhardt J, Radke J, Dittmayer C, Franz J, Thomas C, Mothes R, Laue M, Schneider J, Brünink S, Greuel S, Lehmann M, Hassan O, Aschman T, Schumann E, Chua RL, Conrad C, Eils R, Stenzel W, Windgassen M, Rößler L, Goebel HH, Gelderblom HR, Martin H, Nitsche A, Schulz-Schaeffer WJ, Hakroush S, Winkler MS, Tampe B, Scheibe F, Körtvélyessy P, Reinhold D, Siegmund B, Kühl AA, Elezkurtaj S, Horst D, Oesterhelweg L, Tsokos M, Ingold-Heppner B, Stadelmann C, Drosten C, Corman VM, Radbruch H, Heppner FL. 2020. Olfactory transmucosal SARS-CoV-2 invasion as a port of central nervous system entry in individuals with COVID-19. Nat Neurosci https://doi.org/10.1038/s41593-020-00758-5.

40. Hayn M, Hirschenberger M, Koepke L, Straub JH, Nchioua R, Christensen MH, Klute S, Bozzo CP, Aftab W, Zech F, Conzelmann C, Müller JA, Badarinarayan SS, Stürzel CM, Forne I, Stenger S, Conzelmann K-K, Münch J, Sauter D, Schmidt FI, Imhof A, Kirchhoff F, Sparrer KMJ. 2020. Imperfect innate immune antagonism renders SARS-CoV-2 vulnerable towards IFN- γ and λ - . bioRxiv 2020.10.15.340612.

41. Thoms M, Buschauer R, Ameismeier M, Koepke L, Denk T, Hirschenberger M, Kratzat H, Hayn M, Mackens-Kiani T, Cheng J, Straub JH, Stürzel CM, Fröhlich T, Berninghausen O, Becker T, Kirchhoff F, Sparrer KMJ, Beckmann R. 2020. Structural basis for translational shutdown and immune evasion by the Nsp1 protein of SARS-CoV-2. Science (80-) 369:1249 LP – 1255.

42. García-Sastre A. 2017. Ten Strategies of Interferon Evasion by Viruses. Cell Host Microbe 22:176–184.

43. Huang J, Hume AJ, Abo KM, Werder RB, Villacorta-Martin C, Alysandratos KD, Beermann M Lou, Simone-Roach C, Lindstrom-Vautrin J, Olejnik J, Suder EL, Bullitt E, Hinds A, Sharma A, Bosmann M, Wang R, Hawkins F, Burks EJ, Saeed M, Wilson AA, Mühlberger E, Kotton DN. 2020. SARS-CoV-2 Infection of Pluripotent Stem Cell-Derived Human Lung Alveolar Type 2 Cells Elicits a Rapid Epithelial-Intrinsic Inflammatory Response. Cell Stem Cell 962–973.

44. Stanifer ML, Kee C, Cortese M, Zumaran CM, Triana S, Mukenhirn M, Kraeusslich HG, Alexandrov T, Bartenschlager R, Boulant S. 2020. Critical Role of Type III Interferon in Controlling SARS-CoV-2 Infection in Human Intestinal Epithelial Cells. Cell Rep 32.

45. Puelles VG, Fleck D, Ortz L, Papadouri S, Strieder T, Böhner AMC, van der Wolde JW, Vogt M, Saritas T, Kuppe C, Fuss A, Menzel S, Klinkhammer BM, Müller-Newen G, Heymann F, Decker L, Braun F, Kretz O, Huber TB, Susaki EA, Ueda HR, Boor P, Floege J, Kramann R, Kurts C, Bertram JF, Spehr M, Nikolic-Paterson DJ, Moeller MJ. 2019. Novel 3D analysis using optical tissue clearing documents the evolution of murine rapidly progressive glomerulonephritis. Kidney Int 96:505–516.

