## Supplementary figures and images for "Human nasal and lung tissues infected *ex vivo* with SARS-CoV-2 provide insights into differential tissue-specific and virus-specific innate immune responses in the upper and lower respiratory tract"

### Supplemental figure 1

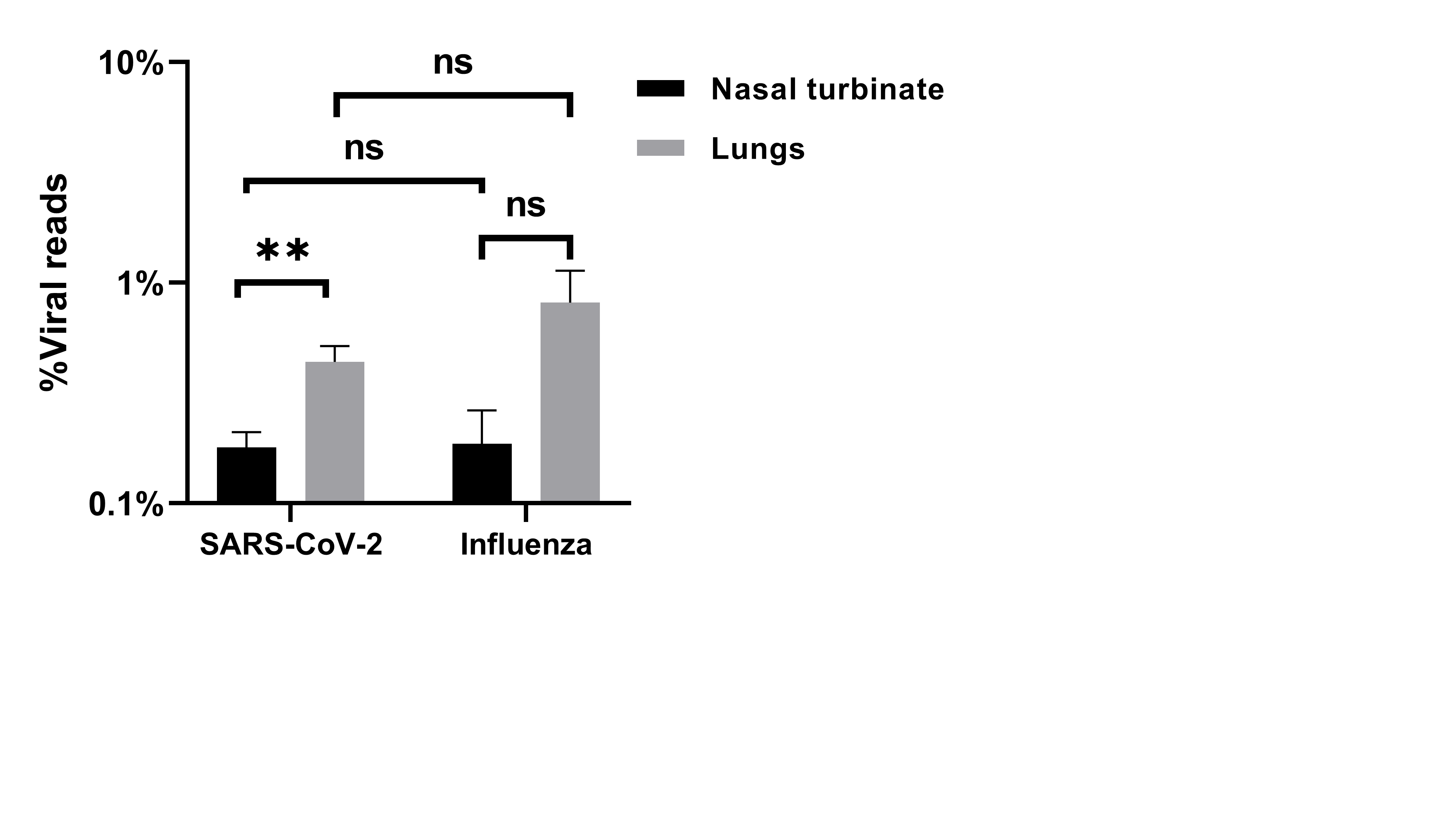

### Supplemental figure 2

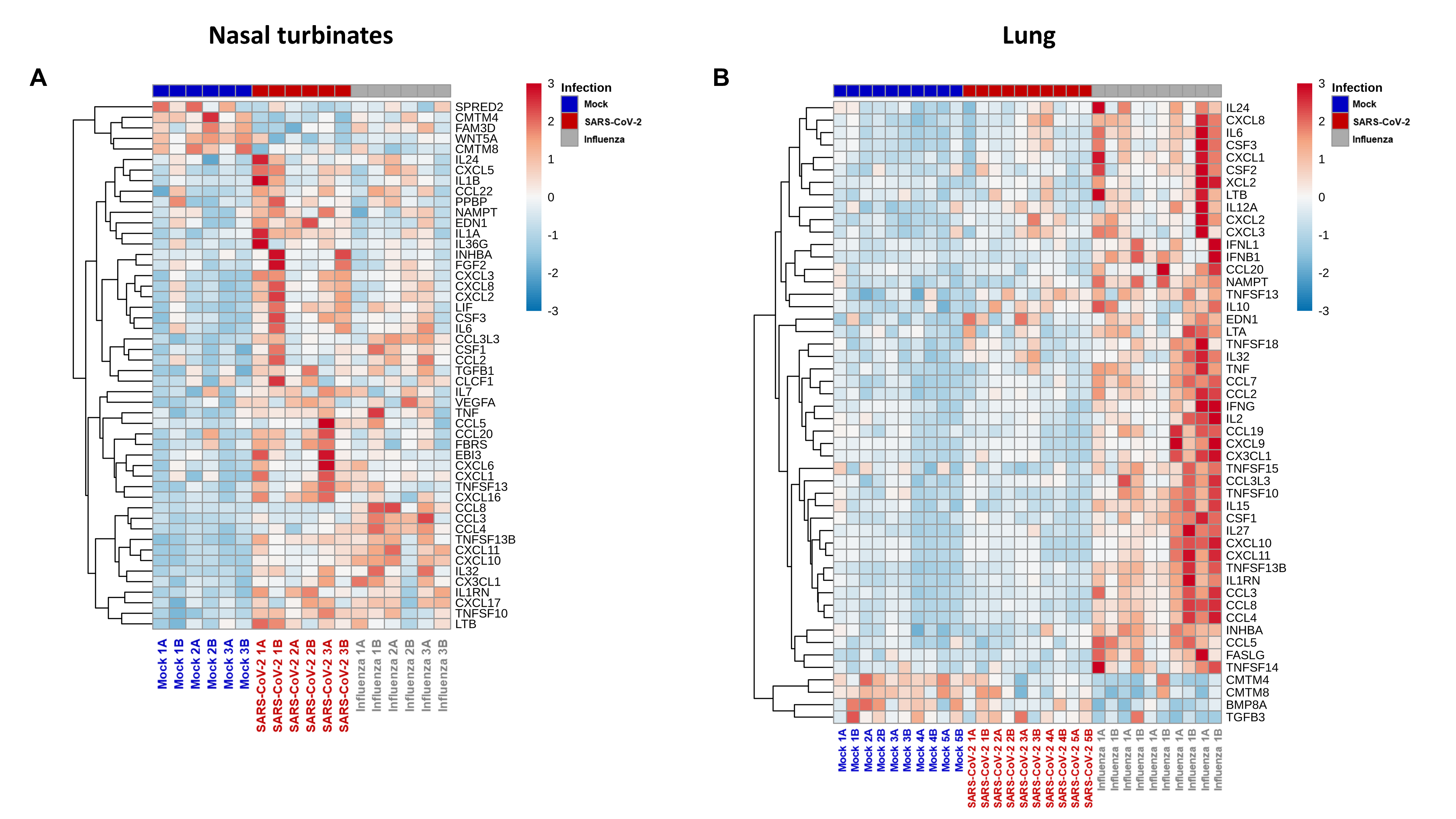

### Supplemental figure 3

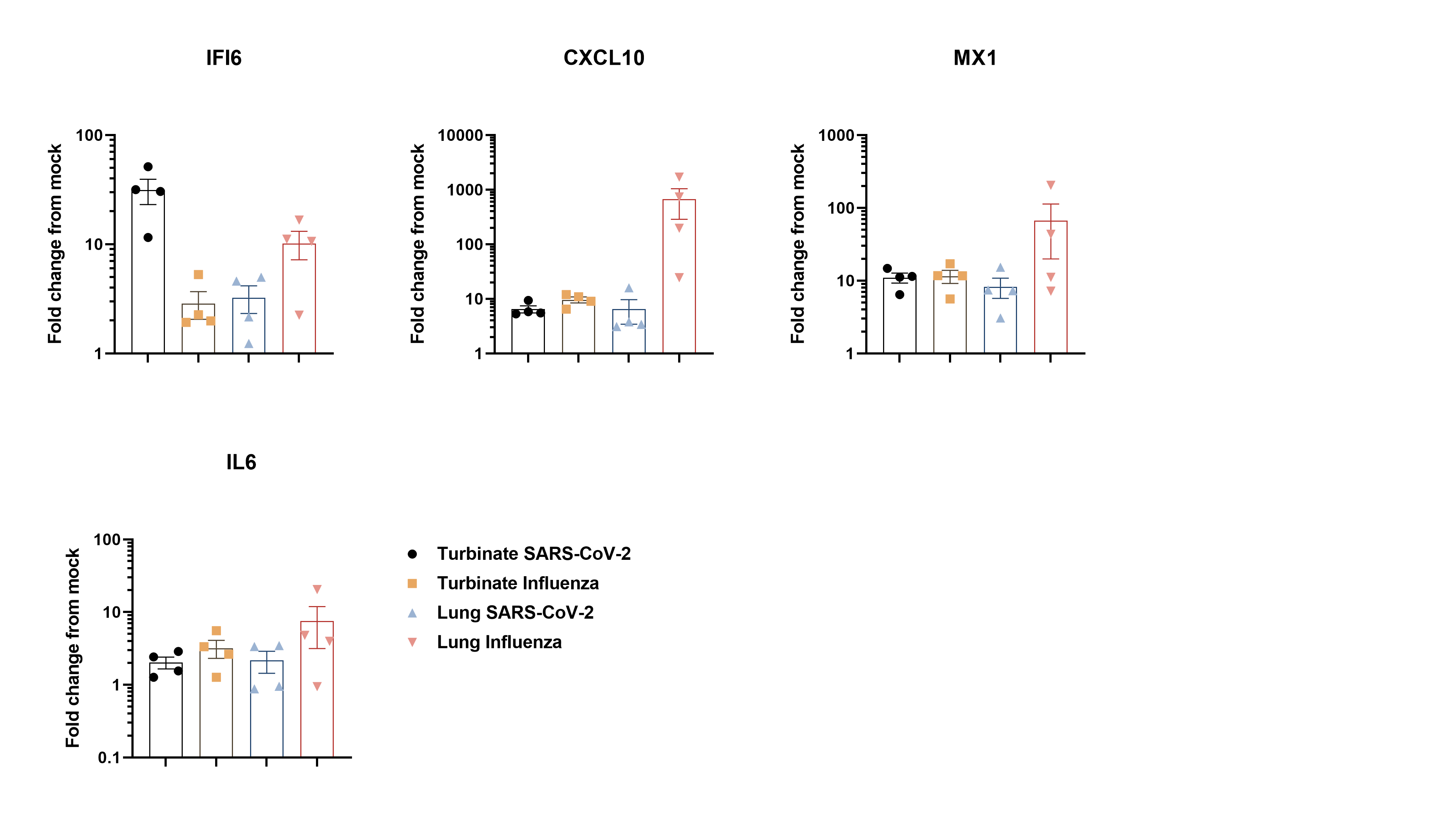
