## Supplemental Table 1 for "Human nasal and lung tissues infected *ex vivo* with SARS-CoV-2 provide insights into differential tissue-specific and virus-specific innate immune responses in the upper and lower respiratory tract"

TABLE S1. Primers and probes for real time PCR analysis

| **Target** | **Sequence** |
| --- | --- |
| **SARS-CoV-2 sub-genomic N gene** | Forward: 5'-CCCAGGTAACAAACCAACCAAC-3' |
|  | Reverse: 5'-AGTGAGAGCGGTGAACCAAG-3' |
|  | Probe: 5'-CGTTTGGTGGACCCTCAGATTCAACTGGCA-3' |
| **SARS-CoV-2 genomic E gene** | Forward: 5'-ACAGGTACGTTAATAGTTAATAGCGT-3' |
|  | Reverse: 5'-ATATTGCAGCAGTACGCACACA-3' |
|  | Probe: 5'-ACACTAGCCATCCTTACTGCGCTTCG-3' |
| **Influenza A/H1N1 M2 gene** | Forward: 5'-GACCRATCCTGTCACCTCTGAC-3' |
|  | Reverse: 5'-AGGGCATTYTGGACAAAKCGTCTA-3' |
|  | Probe: 5'-TGCAGTCCTCGCTCACTGGGCACG-3' |
| **IFNα** | Forward: 5'-GGCTGTGAGGAAATACTTCCAAAGAA-3' |
|  | Reverse: 5'-GATCTCATGATTTCTGCTCTGACAAC-3' |
| **IFNβ** | Forward: 5'- TGGGAGGCTTGAATACTGCCTCAA -3' |
|  | Reverse: 5'- TGCGGCGTCCTCCTTCTGGA -3' |
|  | Probe: 5'-CCTGAGGAGATTAAGCAGCTGCAGC-3' |
| **IFNγ** | Forward: 5'-GCAACAAAAAGAAACGAGATGACTTCG-3' |
|  | Reverse: 5'-TGAGTTCATGTATTGCTTTGCGTTG-3' |
|  | Probe: 5'-AGCTGACTAATTATTCGGTAACTGA-3' |
| **IFNλ1** | Forward: 5'- CGCCTTGGAAGAGTCACTCA -3' |
|  | Reverse: 5'- GAAGCCTCAGGTCCCAATTC -3' |
| **IFNλ2** | Forward: 5'- ACATAGCCCAGTTCAAGTC -3' |
|  | Reverse: 5'- GACTCTTCTAAGGCATCTTTG -3' |
| **MX1** | Forward: 5'-GGAGATCTTTCAGCACCTG-3' |
|  | Reverse: 5'-ACGTCTGGAGCATGAAGAACTG-3' |
|  | Probe: 5'-CCTATCACCAGGAGGCCAGC-3' |
| **CXCL10** | Forward: 5'-GAAAAACTTGAAATTATTCCTGCAAGCC-3' |
|  | Reverse: 5'-TGGATTCAGACATCTCTTCTCACCC-3' |
|  | Probe: 5'-TCCACGTGTTGAGATCATTGCTACAA-3' |
| **IL6** | Forward: 5'-AGAAAACAACCTGAACCTTCC-3' |
|  | Reverse: 5'-ATACCTCAAACTCCAAAAGACC-3' |
| **IFI6** | Forward: 5'-GGTGGAGGCAGGTAAGAAAAAG-3' |
|  | Reverse: 5'-ATCGCAGACCAGCTCATCAG-3' |
| **β-actin** | Forward: 5'-CCTGGCACCCAGCACAAT-3' |
|  | Reverse: 5'- GCCGATCCACACGGAGTACT-3' |
|  | Probe: 5'- ATCAAGATCATTGCTCCTCCTGAGCGC-3' |
